# Pre-diabetes linked microRNA miR-193b-3p targets PPARGC1A and increases lipid accumulation in hepatocytes: potential biomarker of fatty liver disease

**DOI:** 10.1101/2021.01.22.427820

**Authors:** Inês Guerra Mollet, Maria Paula Macedo

## Abstract

Distinct plasma microRNA profiles associate with different disease features. Elevated plasma microRNA hsa-miR-193b-3p has been reported in patients with pre-diabetes where early liver dysmetabolism plays a crucial role. MicroRNA target databases revealed that hsa-miR-193b-3p could potentially target PPARGC1A, a master switch transcriptional co-activator that orchestrates the expression of genes involved in multiple metabolic pathways. In this study we evaluated the effects of hsa-miR-193b-3p in the human HepG2 hepatocyte cell line. We show that hsa-miR-193b-3p targets PPARGC1A/PGC1α mRNA 3’UTR and reduces its expression. Overexpression of hsa-miR-193b-3p under hyperglycemia resulted in increased accumulation of intracellular lipid droplets; and quantitative real-time RT-PCR of a panel of 24 genes revealed significant disruption in the expression of genes coordinating glucose uptake, glycolysis, pyruvate metabolism, fatty acid synthesis, fatty acid oxidation, triglyceride synthesis, VLDL secretion, insulin signalling, and mitochondrial biogenesis, dynamics and function. These results provide *in vitro* evidence that microRNA hsa-miR-193b-3p downregulates PPARGC1A/PGC1α, disrupts downstream metabolic pathways and leads to the accumulation of intracellular lipids in hepatocytes. We propose that microRNA hsa-miR-193b-3p may have potential as a clinically relevant plasma biomarker for metabolic associated fatty liver disease in a pre-diabetic dysglycemic context.

## Introduction

MicroRNAs are small non-protein coding 20-22 nucleotide long RNAs produced by cells of various tissues that can be actively released into the bloodstream in vesicles called exosomes and delivered to cells in the same or other tissues in the organism (1). MicroRNAs target protein expression essentially by binding to the 3’ untranslated region (3’UTR) of messenger RNA (mRNA) thereby preventing translation of the mRNA into protein and directing the mRNA to nonsense mediated decay (2). We hypothesize that particular microRNAs detected in circulation in early disease states can be used as biomarkers of underlying dysmetabolism and also that the targets (3) of each microRNA can provide specific crucial organ specific pathological information that could be used to inform early clinical intervention. Several circulating miRNAs have been reported as being altered in plasma from patients with metabolic disease including pre-diabetes, however, the specific mechanisms through which they function remain unexplored (4).

Pre-diabetes, diagnosed as impaired fasting glucose and/or impaired glucose tolerance, is considered an early sign of metabolic syndrome (5). NAFLD/MAFLD (Nonalcoholic Fatty Liver Disease/Metabolic Associated Fatty Liver Disease) (6) is a precursor of metabolic syndrome (7, 8). NAFLD/MAFLD is a condition characterized by a fatty liver, which can originate complications, including cirrhosis, liver failure, liver cancer, and cardiometabolic health problems that encompass dysmetabolism. Thus, analyzing the effects of pre-diabetes associated circulating microRNA targets in liver cells is an appropriate premise to identify early intervention targets that may have potentially important clinical application in diagnosis, prevention and treatment, paving the way for precision medicine. Among these is microRNA hsa-miR-193b-3p, which is overexpressed in plasma of patients with impaired fasting glucose and glucose intolerance and which tends to normalize upon exercise intervention in humans (4, 9) but for which no data exists on a precise mechanism or relevant targeting.

In this study we demonstrate that hsa-miR-193b-3p targets the expression of PPARGC1A mRNA and increases lipid accumulation in human hepatocyte derived cells. PPARGC1A codes for PGC-1α a central co-activator of transcriptional cascades that regulate several interconnected pathways including glucose metabolism (10), lipid metabolism (11, 12), inflammation, mitochondrial function and oxidative stress (13). PGC-1α functions by interacting with the nuclear peroxisome proliferator-activated receptors (PPARs) activating them to bind DNA in complex with retinoid X receptor (RXR) (14), on the promoter of numerous genes central to metabolic regulation, increasing transcription of specific genes and decreasing transcription of others. The peroxisome proliferator activated receptors (PPARs) are nuclear receptors that play key roles in the regulation of lipid and glucose metabolism, inflammation, cellular growth, and differentiation. The receptors bind and are activated by a broad range of fatty acids and fatty acid derivatives serving as major transcriptional sensors of fatty acids (14). We therefore also analyzed the expression of several metabolic master-switch genes downstream of hsa-miR-193b-3p downregulation of PPRAGC1A/PGC1α to analyze downstream pathway regulation. Glucose and lipid metabolism depend on the ability of mitochondria to generate energy in cells (15), therefore we also looked at hepatic mitochondrial dysfunction which plays a central role in the pathogenesis of insulin resistance in obesity, pre-diabetes and NAFLD/MAFLD (16).

We sought to investigate whether elevated levels of microRNA hsa-miR-193b-3p in hepatocytes could interfere with hepatic cell function *in vitro* under conditions that mimic pre-diabetes. We evaluated pre-diabetes associated microRNA hsa-miR-193b-3p predicted target PPARGC1A (PGC1α), hepatic lipid content and direct or indirect effects on expression of genes regulating cellular energy metabolism, in human hepatocyte derived cells. The objective was to investigate which metabolic pathways microRNA hsa-miR-193b-3 interfered with so as to gage the potential of hsa-miR-193b-3p as an early plasma biomarker of specific liver dysmetabolism to support precision medicine in early stages.

## Results

### MicroRNA hsa-miR-193b-3p targets the 3’UTR of PPARGC1A mRNA and downregulates PPARGC1A expression in HepG2 cells

To evaluate potential mRNA targets of microRNA hsa-miR-193b-3p we performed a bioinformatic analysis of conserved predicted targets of hsa-miR-193b-3p from the online database TargetScan 7.1 (3) using the DAVID Bioinformatics Resources 6.8 (17, 18) and clustering tools for functional Gene Ontology Annotations (19). Of the 283 predicted conserved mRNA targets of microRNA hsa-miR-193b-3p we found the top gene ontology cluster was composed of 43 genes involved in gene transcription. From this list PPRAGC1A stood out as the gene with the most consistently conserved microRNA hsa-miR-193b-3p target site involved in energy metabolism. Peroxisome Proliferator-Activated Receptor Gamma Coactivator 1-Alpha (PPARGC1A) codes for PGC-1α, a transcriptional coactivator of numerous genes involved in energy metabolism. When hsa-miR-193b-3p was overexpressed in HepG2 cells for 72 hours using a mimic, we observed robust downregulation of PPARGC1A mRNA expression at basal glucose concentration (5mM glucose, Figure 1A), under hyperglycemia (20 mM glucose, Figure 1B), under hyperinsulinemia (5mM glucose and final 24h 10nM insulin, Figure 1C) and under hyperglycemia/hyperinsulinemia (20mM glucose and final 24h 10nM insulin, Figure 1D). We validated the predicted PPARGC1A 3’UTR target site for miR-193b-3p by cloning a “match”, or a “mismatch” control (Figure 1E), into the 3’UTR of the firefly luciferase gene in the pmirGLO Dual-Luciferase miRNA Target Expression Vector (Figure 1F), followed by dual-glow luciferase assay. Expression of firefly luciferase from the “match” clone was significantly downregulated in HepG2 cells when the hsa-miR-193b-3p mimic was overexpressed and cells were transfected with pmirGLO match clone (Figure 1G). These results support our posit that microRNA hsa-miR-193b-3p directly targets and downregulates the human PPARGC1A mRNA by binding to its 3’UTR.

**Figure 1.**
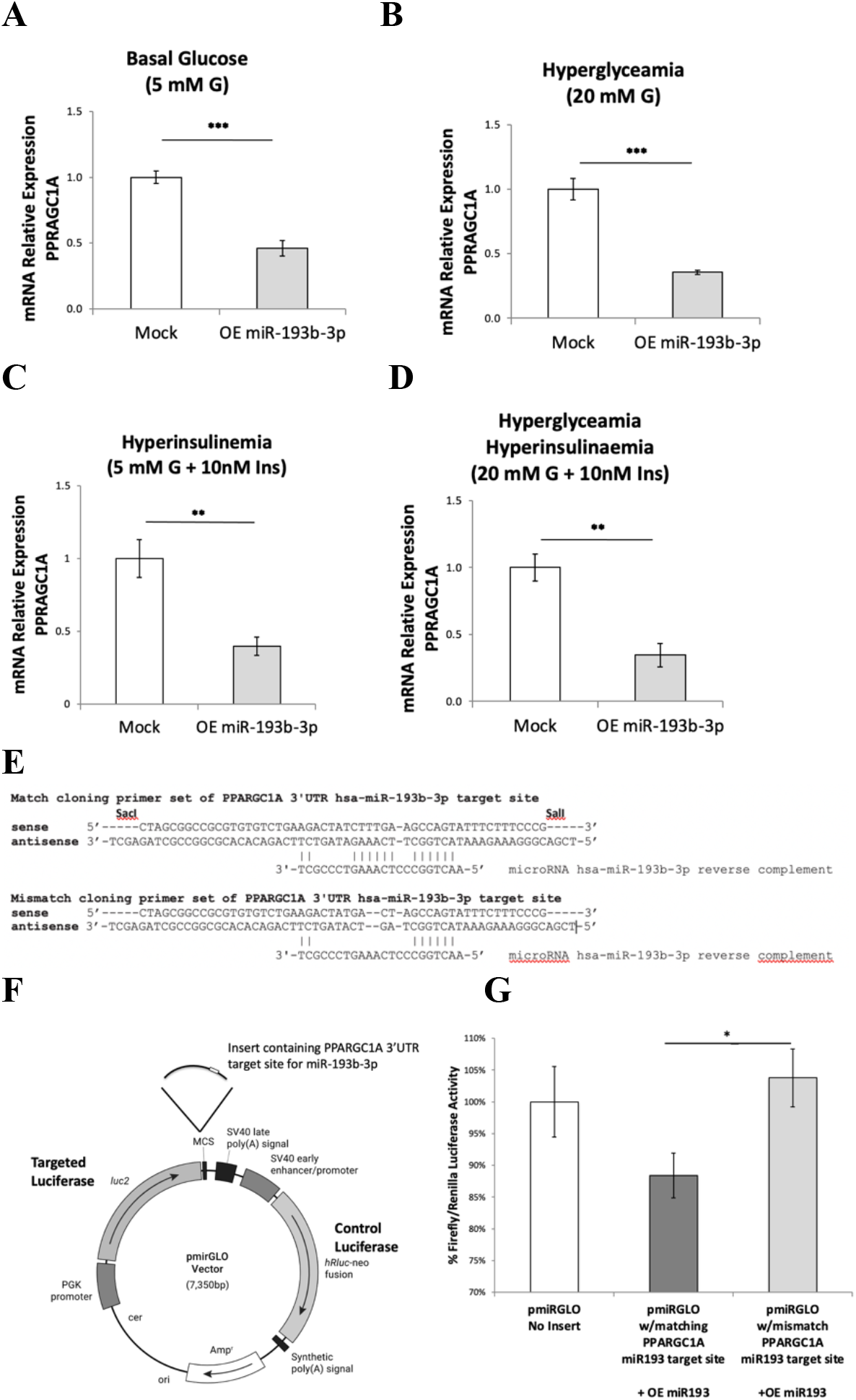
MicroRNA hsa-miR-193b-3p targets the 3’UTR of PPARGC1A mRNA and downregulates PPARGC1A expression in HepG2 cells. *(A, B, C, D)* mRNA expression, evaluated by real-time RT-qPCR, of PPARGC1A after 72h overexpression of hsa-miR-193b-3p in HepG2 cells at basal 5 mM glucose *(A)*, under hyperglycemia at 20 mM glucose *(B)*, under hyperinsulinemia with 5 mM glucose and final 24 hours with 10nM insulin *(C)* and hyperglycemia /hyperinsulinemia at 20mM glucose and final 24hours with 10nM insulin *(D)*. *(E)* Match and mismatch primer pairs containing the PPARGC1A 3’UTR target site for hsa-miR-193b-3p that were cloned into pmirGLO miRNA target expression vector (*F)*. *(G)* Luciferase luminescence measurement after overexpression of hsa-miR-193b-3p in HepG2 cells; pmirGLO plasmid was transfected into HepG2 cells 24h before luciferase assay. Data are presented as mean ± SEM of n=4. Mock - mock transfection, OE - overexpression of hsa-miR-193b-3p, G - glucose, Ins - insulin. Student’s t-test p-values indicated by * < 0.05, ** < 0.01, *** < 0.001.

### Intracellular lipid droplet content is increased by overexpression of microRNA hsa-miR-193b-3p in HepG2 cells

PPARGC1A/PGC1α is of central importance for lipid regulation (11). Fatty liver is associated with impaired activity of PPARGC1A (20). Therefore, having verified that hsa-miR-193b-3p overexpression in HepG2 hepatoma cells reduced the expression of PPARGC1A we proceeded to analyze the intracellular lipid content in these cells. We overexpressed hsa-miR-193b-3p mimic in HepG2 cells for 72h, under normoglycemia (5mM glucose) and hyperglycemia (20mM glucose), visualized lipid droplet content using Oil Red O staining and brightfield microscopy (Figure 2) and performed quantitative image analysis. At hyperglycemia (20mM glucose) we observed a significant increase in total area covered by lipid droplets (Figure 2G) and number of lipid droplets (Figure 2H); we also saw an increasing trend in lipid droplet size (Figure 2I), though not statistically significant. All three measurements also showed increasing trends at 5mM (data not shown). This result supports further research into the possibility of using increased plasma levels of microRNA hsa-miR-193b-3p as a biomarker for fatty liver.

**Figure 2.**
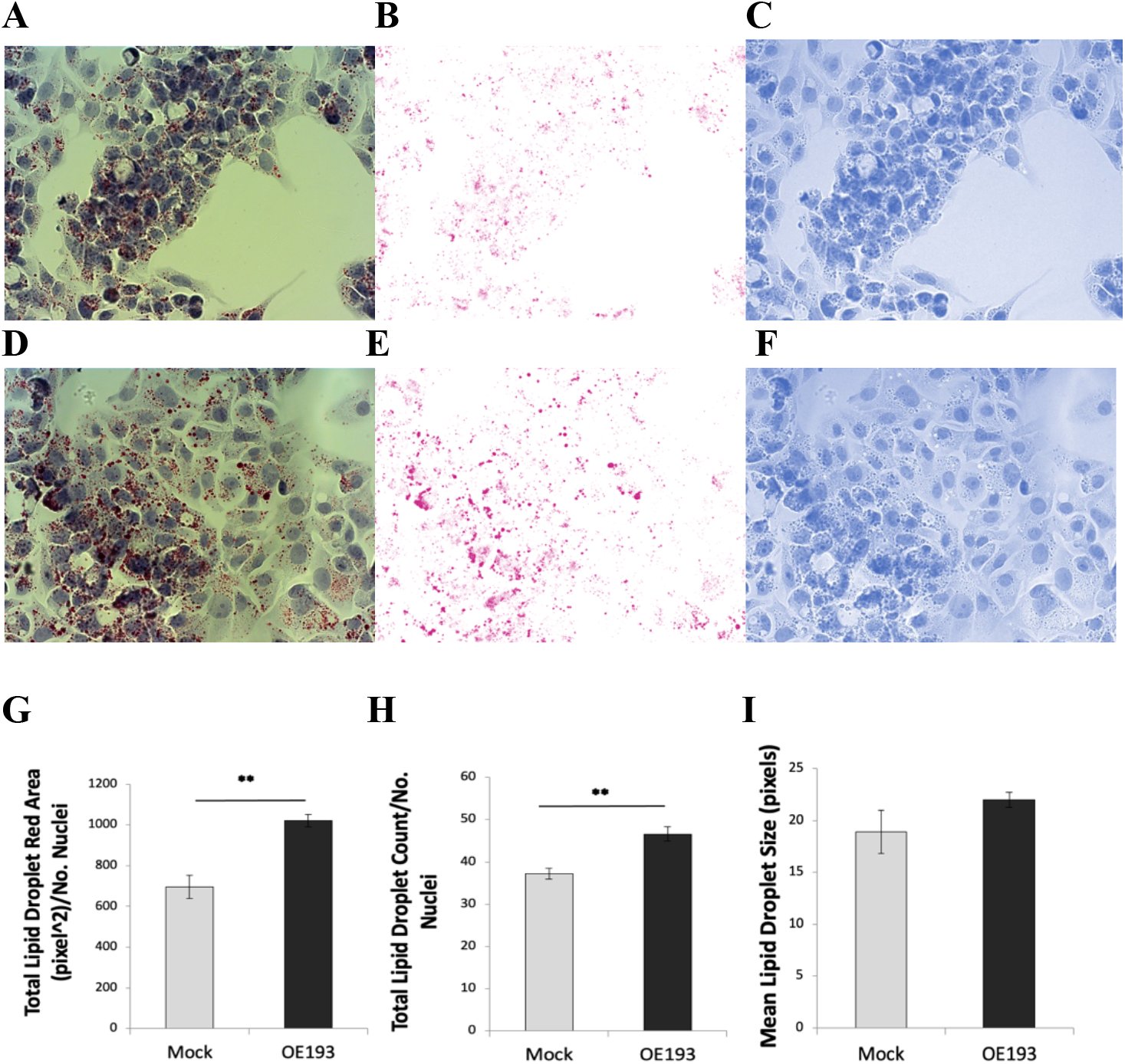
MicroRNA hsa-miR-193b-3p overexpression in HepG2 cells increases intracellular lipid droplet accumulation. HepG2 cells cultured at 20mM glucose. Lipid droplets in HepG2 cells were visualized using Oil Red O staining. Brightfield 400x magnification images of Oil Red O staining in control mock transfection (A) with corresponding red (B) and blue (C) color deconvolution. Brightfield 400x magnification images of Oil Red O staining after microRNA hsa-miR193b-3p overexpression (D) with corresponding red (E) and blue (F) color deconvolution. Red color deconvolution used for red particle analysis; blue color deconvolution used for nuclei counting. Quantitative analysis of lipid droplet area (G), lipid droplet number (H) and lipid droplet size (I). Data are presented as mean ± SEM of n=4; each n is average of with ten fields per condition normalized to number of nuclei. Mock - mock transfection, OE - overexpression of hsa-miR-193b-3p. Student’s t-test p-value ** < 0.01.

### Changes in expression of genes involved in lipid metabolism may in part explain intracellular lipid accumulation following overexpression of hsa-miR-193b-3p in HepG2 cells

Having observed that PPARGC1A expression is robustly downregulated and lipid droplets accumulate when microRNA hsa-miR-193b-3p is overexpressed in HepG2 cells under hyperglycemia-hyperinsulinemia we proceeded to analyze the expression of other genes involved in lipid metabolism and processing that might be affected. The lower mRNA expression of Microsomal Triglyceride Transfer Protein, MTTP (Figure 3A) which is required for the secretion of plasma lipoproteins containing apolipoprotein B (21), such as VLDL, that is made in hepatocytes and secreted to deliver lipids, may be in part responsible for the increased lipid droplet accumulation observed. The increased expression of Tribbles Pseudokinase 1, TRIB1 (Figure 3B), could also be responsible for the lipid droplet accumulation we observed (22); indeed Burkhardt et al. (2010) reported that hepatic overexpression of TRIB1 in mouse led to reduce VLDL secretion. The increased expression of Low Density Lipoprotein Receptor, LDLR (Figure 3C), would support increased LDL uptake in favor of increased lipid accumulation. No significant expression changes were observed in Apolipoprotein B, APOB (Figure 3D).

**Figure 3.**
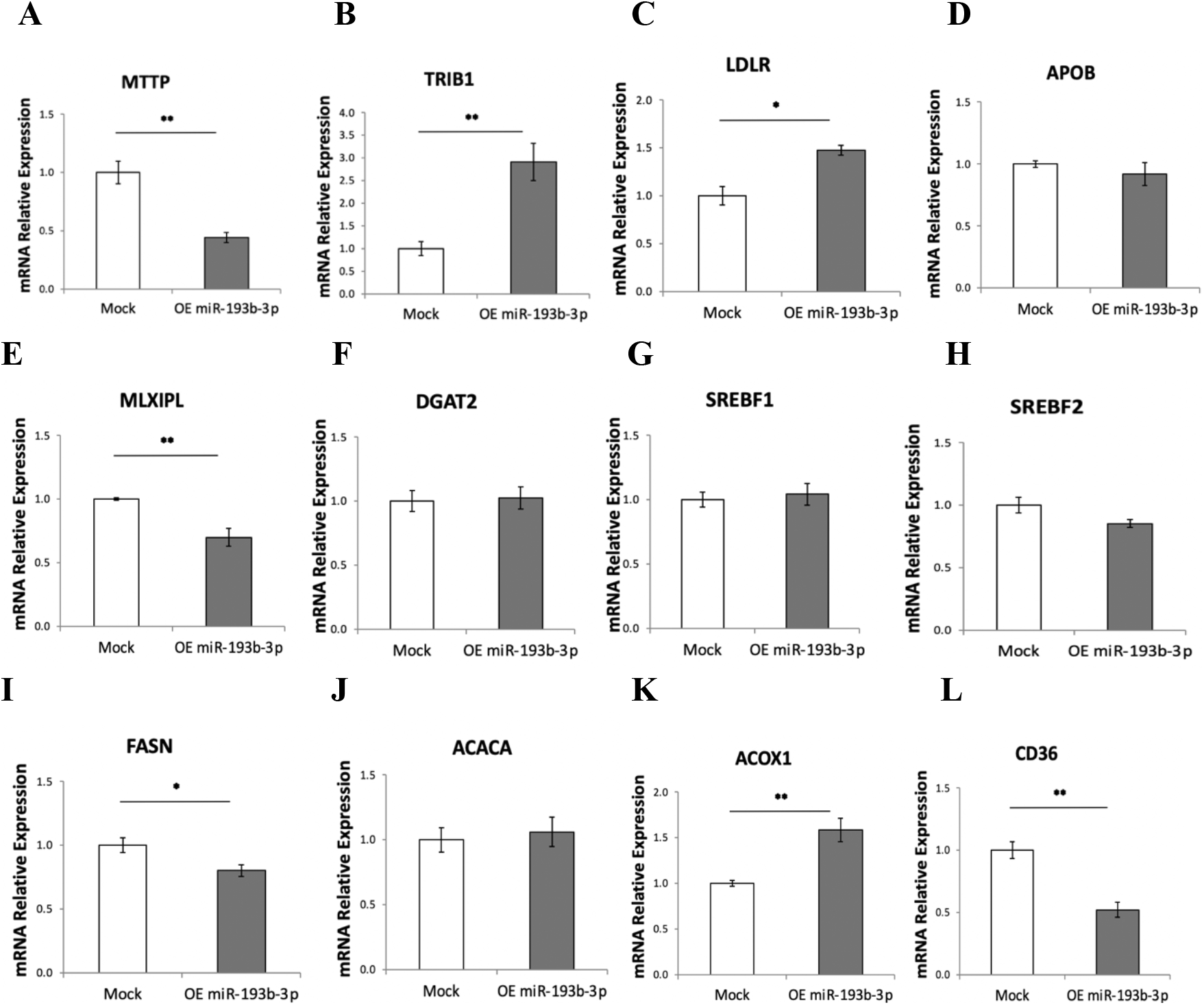
Overexpression of microRNA hsa-miR-193b-3p in HepG2 alters mRNA expression of fundamental genes involved in lipid processing. Lower mRNA expression of MTTP (A) and higher expression of TRIB1 (B) suggest reduced VLDL secretion. Higher level of LDL Receptor LDLR (C, n=3). No change in APOB (D). Lower mRNA levels of and MLXIPL/ChREBP (E, Mock n=4, OE n=3) coordinating triglyceride synthesis. No change in DGAT2 (F), SREBF1 (G, n=3) or SREBF2 (H, n=3). Lower expression of fatty acid synthase FASN (I). No change in ACACA (J). Increased ACOX1 (K) indicates increased fatty acid oxidation. Reduced fatty acid translocase CD36 (L). Relative mRNA expression was evaluated by real-time RT-qPCR after 72h overexpression of hsa-miR-193b-3p in HepG2 cells under hyperglycemia/hyperinsulinemia (20mM glucose, with 10nM insulin during last 24 hours). Data are presented as mean ± SEM of n=4, unless otherwise indicated. Mock - mock transfection, OE - overexpression of hsa-miR-193b-3p. Student’s t-test p-values indicated by * < 0.05, ** < 0.01, *** < 0.001.

Expression of carbohydrate-responsive element-binding protein (ChREBP) also known as MLX-interacting protein-like, MLXIPL, was reduced by 30% (Figure 3E) when hsa-miR-193b-3p was overexpressed. This transcription factor binds and activates carbohydrate response element (ChoRE) motifs on the promoters of triglyceride synthesis genes in a glucose-dependent manner. Thus we would expect to see reduced expression of genes involved in triglyceride synthesis such as DGAT2 (23), which catalyzes the final reaction in triglyceride synthesis. However, no changes in expression of DGAT2 (Figure 3F) were observed. No expression changes were observed in either of two major transcriptional regulators, Sterol Regulatory Element Binding Transcription Factors 1 and 2 coded by SREBF1 (Figure 3G) and SREBF2 (Figure 3H), which in concert with ChREBP regulate hepatic lipid metabolism by inducing transcription of lipogenic enzymes that direct lipogenesis in the liver (24). We also observed a 20% reduction in expression of Fatty Acid Synthase, FASN (Figure 3I) indicating reduced fatty acid synthesis, which was expected since this gene is regulated by ChREBP/MLXIPL. However, no change was observed in Acetyl-CoA carboxylase, ACACA (Figure 3J), the enzyme which catalyzes the carboxylation of acetyl-CoA to malonyl-CoA, the rate-limiting step in fatty acid synthesis. This lack of downregulation of ACACA is important, because although we observe increased expression of Acyl-CoA Oxidase 1, ACOX1 (Figure 3K) indicating increased fatty acid oxidation in the mitochondria, the malonyl-CoA produced by ACACA in the cytoplasm would be expected inhibit transfer of cytoplasmic fatty acids into the mitochondria for beta oxidation which would contributing to increased lipids in the cytoplasm. Unexpectedly, we observed a 50% reduction in the fatty acid translocate CD36 (Figure 3L), which has been described as positively correlated with fatty liver (25).

### Changes favoring insulin signaling, reduced glycolysis, and reduced mitochondrial activity are observed when microRNA hsa-miR-193b-3p is overexpressed in HepG2 cells

In addition to lipid metabolism PPARGC1A/PGC1α also regulates insulin signaling, hepatic glucose metabolism and mitochondrial turnover (20, 26, 27). Therefore, we investigated whether expression of genes involved in these mechanisms were altered. We observed downregulation of YWHAZ (Figure 4A). YWHAZ belongs to the 14-3-3 family of proteins that mediate signal transduction; it binds to insulin receptor substrate-1 (IRS-1) and is thought to interrupt the association between the insulin receptor and IRS-1 thus interfering with insulin signaling (28). Our result indicates decreased interference with insulin signaling by this particular mechanism. No change was observed in Tribbles homolog 3, TRIB3 (Figure 4B) that has been linked to insulin resistance and hepatic production of glucose in the liver (29, 30).

**Figure 4.**
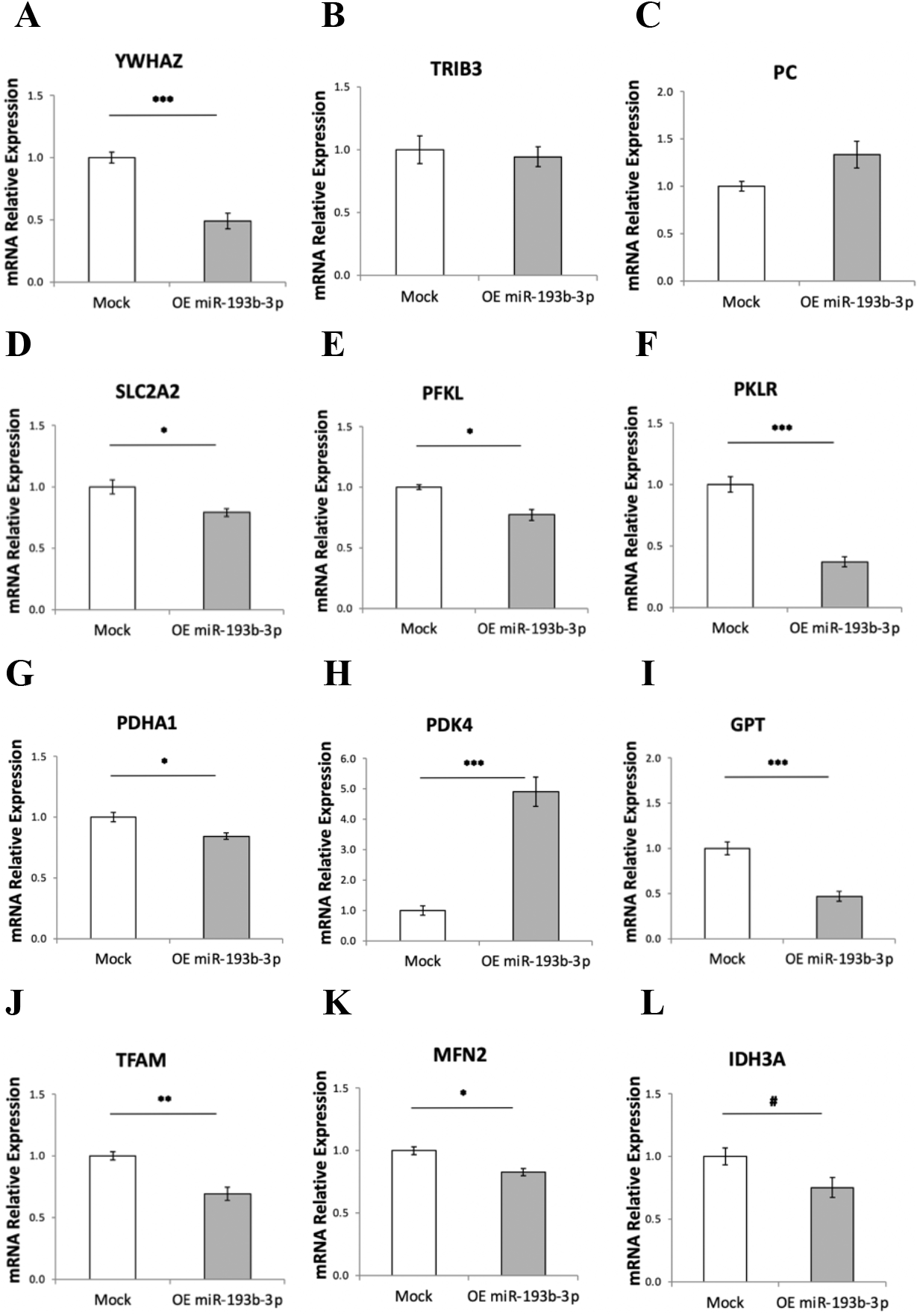
Changes in genes expression regulating of insulin signaling, glucose metabolism, pyruvate availability and mitochondrial activity when hsa-miR-193b-3p is overexpressed in HepG2 cells. Downregulation of YWHAZ (A) indicates increased insulin signaling. No change in TRIB3 (B, n=3) linked to insulin resistance. No change in pyruvate carboxylase, PC (C, n=3). Reduced glucose transporter SLC2A2/GLUT2 (D, n=3) indicates reduced glucose transport. Reduced levels of phosphofructokinase PFKL (E, n=3) and pyruvate kinase PKLR (F) indicate reduced glycolysis. Reduced levels of pyruvate dehydrogenase PDHA1 (G) and increase in its inhibiting kinase PDK4 (H) indicate lower conversion of pyruvate to Acetyl-CoA. Reduced Glutamic-Pyruvic Transaminase GTP (I) indicates lower conversion of alanine to pyruvate. Reduced levels of mitochondrial Transcription Factor A, TFAM (J), mitofusin 2, MFN2 (K, Mock n=4, OE193 n=3) and Isocitrate Dehydrogenase 3 Catalytic Subunit Alpha IDH3A (L, n=3) indicate reduced levels of mitochondrial biogenesis, fusing dynamics and function respectively. Relative mRNA expression was evaluated by real-time RT-qPCR after 72h overexpression of hsa-miR-193b-3p in HepG2 cells under hyperglycemia/hyperinsulinemia (20mM glucose, with 10nM insulin during last 24 hours). Data are presented as mean ± SEM of n=4, unless otherwise indicated. Mock - mock transfection, OE - overexpression of hsa-miR-193b-3p. Student’s t-test p-values indicated by #<0.01, * < 0.05, ** < 0.01, *** < 0.001.

PPARGC1A/PGC1α is known to regulate key hepatic gluconeogenic enzymes leading to increased glucose output (27), however we saw no significant change in expression of pyruvate carboxylase, PC (Figure 4C), the mitochondrial matrix enzyme that regulates fuel partitioning toward gluconeogenesis in hepatocytes. We saw reduced expression of the plasma membrane bidirectional glucose transporter SLC2A2/GLUT2 (Figure 4D) indicating both reduced uptake and release of glucose. The reduced expression of liver phosphofructokinase PFKL (Figure 4E) and pyruvate kinase PKLR (Figure 4F) indicate reduced glycolysis. Reduced expression of the Pyruvate Dehydrogenase E1 Subunit Alpha 1, PDHA1 (Figure 4G), a central component of the pyruvate dehydrogenase complex that is the primary link between glycolysis and the TCA cycle indicated lower conversion of pyruvate to Acetyl-CoA in mitochondria. This is further supported by a five-fold increased expression of the pyruvate dehydrogenase inhibitor mitochondrial kinase PDK4 (Figure 4H). Reduced expression of Glutamic-Pyruvic Transaminase GTP (Figure 4I) indicated lower conversion of alanine to pyruvate. Based on this we could predict a drop in ATP production.

We also observed reduced expression of mitochondrial Transcription Factor A, TFAM (Figure 4J) that is required for mitochondrial biogenesis through maintenance of mitochondrial DNA and replication of normal levels of mitochondrial DNA; reduced expression of mitofusin-2, MFN2 (Figure 4K), that regulates mitochondrial fusion dynamics; and reduced expression of Isocitrate Dehydrogenase 3 Catalytic Subunit Alpha IDH3A (Figure 4L), that catalyzes the rate-limiting step of the TCA cycle in the mitochondrial matrix.

## Discussion

This study is based on recent research showing that circulating microRNAs detected in plasma can enter cells in various tissues and there interfere with gene expression (31), and that, in humans, altered levels of circulating microRNAs are detected in many disease states including metabolic diseases (4). In humans, higher than normal levels of the microRNA hsa-miR-193b-3p have been detected in plasma of patients with impaired fasting glucose and glucose intolerance, two parameters that are used to diagnose pre-diabetes, an early sign of metabolic syndrome (5). Nonalcoholic fatty liver disease, also referred to as metabolic associated fatty liver disease (NAFLD/MAFLD) (6–8), characterized by excess accumulation of lipids in the liver (known as steatosis) is considered a precursor of metabolic syndrome. In this study we have focused on identifying and validating a direct target of hsa-miR-193b-3p and evaluating gene expression changes in cellular energy metabolism driven by the human microRNA hsa-miR-193b-3p in the hepatocyte cell line HepG2, which is considered an appropriate cell model (32). This strategy is in line with recent attempts to try to understand metabolism in type 2 diabetes beyond glycemia (33).

Our results show that microRNA hsa-miR-193b-3p directly targets PPRAGC1A/ PGC1α reducing its expression and that it causes excess intracellular lipid droplet accumulation in HepG2 cells. We also show that the accumulation of lipids observed may be caused in part by altered expression of genes involved in lipid processing which may cause reduced VLDL secretion, as well as reduced glycolysis and reduced mitochondrial activity shifting the metabolic balance toward lipogenesis. We saw a 60% reduction in MTTP required for VLDL secretion, and a 3-fold increased expression of TRIB1 which reduces secretion of VLDL. Indeed, hepatic overexpression of Trib1 in mice reduces secretion of VLDL from liver into circulation (34). Although no change in APOB was observed, the decreased expression levels of MLXIPL/ChREBP suggests a reduced rate of triglyceride synthesis that could lead to insufficient APOB lipidation and secretion.

Insulin signaling normally directs *de novo* lipogenesis in the liver (35), among several other pathways. The observed downregulation of YWHAZ, known to bind insulin receptor substrate-1 (IRS-1) (28) would indicate reduced interference of YWHAZ with IRS-1 at this point in the insulin signaling cascade. However, linking this observation to increased lipogenesis would require investigating additional downstream step in the insulin signaling cascade towards lipogenesis.

When insulin and glucose are associated in liver cells, as in our experimental conditions, the stimulatory effect of glucose on GLUT2 gene expression is predominant (36). However, when hsa-miR-193b-3p is overexpressed we observed reduced SLC2A2/GLUT2, suggesting lower glucose uptake, which would give rise to hyperglycemia, a feature associated with the plasma increase in hsa-miR-193b-3p. We also observed lower glycolysis (reduced PFKL and PKLR) producing less pyruvate; less pyruvate coming from amino acid metabolism (reduced GPT); inhibition of conversion of pyruvate to acetyl-CoA for the TCA cycle (reduced PDHA1 and increased PDK4); all of which would predict a decrease in energy combustion and lower mitochondrial generated ATP. On top of this we observed lower mitochondrial turnover indicated by lower biogenesis, fusion dynamics and activity (reduced TFAM, MFN2 and IDH3A), which would be expected to further contribute to a decrease in energy combustion. In line with our results, mitochondrial dysfunction has been shown to play a central role in the pathogenesis of metabolic diseases and associated complications and reduced energy combustion through reduced glycolysis has been described as critical to excess lipid storage in the liver (37). Energy combustion in the liver is modulated by PPARα-regulated fatty acid beta-oxidation. Interference with PPARα lipid sensing can reduce energy burning and result in accumulation of lipids in hepatocytes. Fatty liver is also associated with impaired activity of PPARGC1A (PGC1α) and reduced mitochondrial biogenesis in mice (20). Although we saw increased expression of ACOX1 which would indicate increased fatty acid beta-oxidation in mitochondria this could be a compensation for reduced mitochondrial biogenesis and dynamics. We saw no change in Acetyl-CoA carboxylase (ACACA), the enzyme which catalyzes the carboxylation of acetyl-CoA to malonyl-CoA the rate-limiting step in fatty acid synthesis; this is relevant, as the malonyl-CoA produced by ongoing fatty acid synthesis will inhibit fatty acid transport into mitochondria for beta-oxidation.

Hepatic disruption of fatty acid translocate CD36 in JAK2L livers has been shown to lower triglyceride (TG), diacylglycerol (DAG) and cholesterol ester (CE) content significantly improving steatosis (25). Therefore, one might speculate that our unexpected observation of a 50% reduction in the fatty acid translocate CD36 alongside lipid accumulation in HepG2 cells under hyperglycemia/ hyperinsulinemia might be a compensatory mechanism.

PPRAGC1A/PGC-1α interacts with the PPAR nuclear receptors (14), on the promoter of numerous genes involved in metabolic regulation. However, the gene expression changes that we observed when microRNA hsa-miR-193b-3p was overexpressed under hyperglycemia/hyperinsulinemia in the HepG2 cell model may result from either direct transcriptional regulation via PPRAGC1A/PGC-1α or other indirect regulation. In future work, one might consider rescuing the downregulation of PPRAGC1A/PGC-1α observed by overexpressing PPRAGC1A/PGC-1α along with hsa-miR-193b-3p in this cell model. As there are a further 283 predicted conserved mRNA targets of microRNA hsa-miR-193b-3p, including 43 genes involved in gene transcription, other predicted gene targets of hsa-miR-193b-3p would also have to be evaluated. With regards to the specific targeting of PPARGC1A/PGC-1α by hsa-miR-193b-3p, and given the central importance of PGC1α in inflammation, oxidative stress and energy in cells other than hepatocytes, much work remains to be done on these pathways and on other cell types and in vivo.

Given that PPRAGC1A/PGC-1α interacts with the nuclear receptor PPAR-γ (14) our results also warrant bearing in mind the drug class of thiazolidinediones, which are potent PPAR-γ agonists, that can be used in the treatment of type 2 diabetes to improve hepatic sensitivity to insulin (38, 39) and improve fibrosis in nonalcoholic steatohepatitis (39, 40). In this context, further work would be required to differentiate effects which are mediated directly through hsa-miR-193b-3p targeting of PPRAGC1A/PGC-1α and PPAR transcriptional regulation, and which might be indirect.

Our results strongly suggest that overexpression of microRNA hsa-miR-193b-3p in hepatoma cells causes a cascade of gene expression changes leading to increased lipid accumulation in these cells, likely triggered in part by direct targeting of PPARGC1A/PGC1α. As levels of plasma hsa-miR-193b-3p are elevated in pre-diabetes in humans (9), the mechanisms we elucidate herein will support the idea of using hsa-miR-193b-3p as an early biomarker of liver dysmetabolism indicative of lipid accumulation in the liver in pre-diabetes. The potential clinical relevance of these results is highlighted by the recent research showing that improvement of fatty liver disease reduces the risk of type 2 diabetes (41). Thus, plasma levels of hsa-miR-193b-3p could conceivably be used to flag pre-diabetic patient for early intervention to target fatty liver thereby preventing aggravation of pre-diabetes and the development of type 2 diabetes.

## Experimental Procedures

### Cell model

The HegG2 hepatoma cell line (32) was maintained in Dulbecco’s Modified Eagle’s Medium (DMEM 25mM high glucose, 21969-035, Gibco, Life Technologies), 4mM glutamine, 10% heat inactivated fetal bovine serum, and 100 IU/ml penicillin, 100 ug/ml streptomycin. Experiments at basal 5mM glucose were performed with DMEM 4mM Glutamine, 1mM Sodium pyruvate, SH30021.FS, Thermo Scientific) 10% heat inactivated fetal bovine serum, 100 IU/ml penicillin, 100 ug/ml streptomycin.

### Overexpression of miR-193b-3p in HepG2 cells

microRNA miR-193b-3p overexpression in HepG2 cells was performed over 72 hours by reverse transfection on day of plating followed by forward transfection the next day, with an additional day of growth, before RNA extraction, plasmid transfection or fixing cells for imaging. MicroRNA mimic for hsa-miR-193b-3p was miRCURY LNA™ microRNA Mimic (472852-001, Exiqon). Transfection of microRNA mimic into HegG2 cells was performed at 5nM final concentration using Lipofectamine® RNAiMAX Transfection Reagent (13778-100, Life Technologies) with 20 min lipofectamine/mimic complex formation in Opti-MEM® I Reduced Serum Medium (31985-062, Life Technologies) and cells incubated in DMEM culture as described above with no penicillin or streptomycin added. Mock transfection was performed with Lipofectamine alone.

### RNA preparation and relative gene expression

Total RNA was prepared by organic extraction using Trizol Reagent (15596-018, Life Technologies) and chloroform (Sigma). RNA was precipitated in absolute ethanol, and dissolved DNase/RNaseFree Distilled Sterile Water (10977-035, Gibco). RNA concentration was determined using Nanodrop 2000 (Thermo Fisher Scientific). Relative genes expression was determined by two step Reverse Transcription Real Time Quantitative PCR (real-time RT-qPCR) with cDNA prepared from total extracted RNA using High-Capacity cDNA Reverse Transcription Kit (4368814, Applied Biosystems) on a PCR Biorad MyCycler (25°C 10 min, 37°C 120 min, 85°C 5 min). For protein coding genes forward and reverse primers for qPCR for each gene of interest were designed across constitutive exons using Primer Quest (Integrated DNA Technologies, IDT, https://www.idtdna.com/pages/tools/primerques t). The list of primers used is presented in Table 1. The custom designed primers were ordered from Invitrogen. Relative gene expression was determined from cDNA using NZYTaq 2× Green Master Mix (MB03903, NZYtech) on MicroAmp Optical 96-well reaction plates (N8010560, Applied Biosystems). Amplification and Ct values for two technical replicates of each sample were obtained on qPCR LightCycler and integrated software (Roche). Amplicon specificity was verified by High Resolution Melting curve using the LightCycler integrated software. Relative mRNA expression was determined using the Relative Gene Expression Data Using Real-Time Quantitative PCR and the 2^(-ddCt) method (42) using TBP primers as reference gene.

**Table 1.**
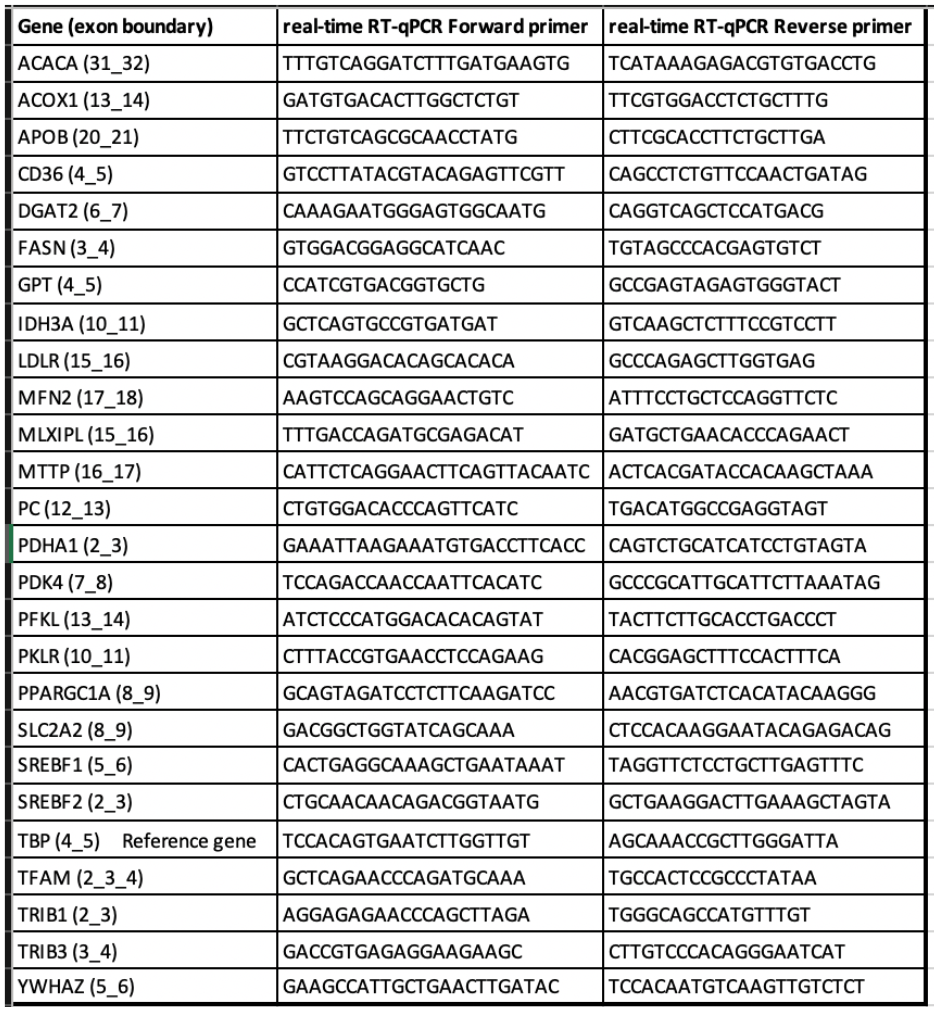
Primers designed for real-time RT-qPCR.

### miRNA target plasmid construct and luciferase assay

To validate the miR-193b-3p target site on the mRNA of PPARGC1A (PGC1a gene), the pmirGLO Dual-Luciferase miRNA Target Expression Vector (E1330, Promega) was used to clone the predicted PPARGC1A target site for microRNA miR-193b-3p along with a mismatch control (Figure 1E,1F). The custom primers for cloning were ordered from Invitrogen. Primer pairs were cloned by double digestion using SacI and SalI restriction enzymes (R0156, R0156, CutSmart Buffer, New England Biolabs) and ligation with T4 DNA Ligase (EL0014, Thermo Fisher Scientific) and FastAP Thermosensitive Alkaline Phosphatase (EF0654, Thermo Fisher Scientific). Selection of clones was made with XhoI/BamHI double digest (R0136, R0146, CutSmart Buffer, New England Biolabs), XhoI site being on the excised multiple cloning site and BamHI being present on the core plasmid. Digestion of positive clones produces a single fragment as opposed to two fragments. Clones were visualized after electrophoresis on 2% agarose using GreenSafe Premium (MB13201, NZYTech) on Chemidoc Touch (Biorad). Plasmids were reproduced by chemical transformation and culture of DH5alpha E. coli (inhouse stock) in LB medium (1% Tryptone, 0.5% Yeast Extract, 1% NaCl, pH 7.0). Colonies were grown on 1.5% Agar/LB (20767.232, VWR Chemicals) with ampicillin selection (20767.232, VWR Chemicals). Plasmid DNA was extracted and purified using Plasmid DNA Mini Kit I (D6943-01, E.Z.N.A.). Plasmid clones were sequenced to verify correct cloned insert with custom sequencing primer 5’-CGAACTGGAGAGCATCCTG-3’ using StabVida SANGER sequencing services (https://www.stabvida.com/sanger-sequencing-service). The selected pmirGLO Dual-Luciferase plasmid clone containing the hsa-miR-193b-3p target site from PPARGC1A was transfected into HepG2 cells grown at 5mM glucose using DharmaFECT kb DNA transfection reagent Lipofectamine (T-2006-01, Dharmacon). Expression of Firefly luciferase was normalized to internal Renilla luciferase using Dual-Glo® Luciferase Assay System (E2920, Promega) with chemiluminescent data obtained in relative light units (RLU) measured using SpectraMax i3x (Molecular Devices).

### Lipid droplet quantification

To identify intracellular lipid content, HepG2 cells were cultured at 5mM glucose or 20mM glucose on 13mm glass coverslips in 24 well plates. At the end of the experiment cells were fixed with 4% paraformaldehyde in phosphate buffered saline at room temperature for 15 min, then washed with phosphate buffered saline before staining with Oil Red O method (43) using a kit (010303, Diapath) according to manufacturer’s instructions. Nuclei were stained blue with hematoxylin. Coverslips were mounted on glass slides for microscopy. Stained cells were analyzed on a Zeiss Z2 fluorescent microscope with ZEN Pro 2012 software. For each condition, ten brightfield color images were captured at 400x magnification with an Axiocam 105 color camera. Image quantitative analysis of lipid droplets was performed on each of ten images per condition with Fiji (ImageJ) software (44) using Color Deconvolution (v1.7, FastRed FastBlue DAB plugin); red color was used for binary/watershed particle analysis to obtain parameters of total red area (pixels^2), average particle size (pixels) and number of particles; blue color was used for manual nuclei counting using Fiji software; number of nuclei per image was used for normalization of red data per image. Results per condition are the average of ten images.

### Statistical Analysis

Statistical analyses were performed using Excel functions. Data is presented as mean ± standard error of the mean (SEM). Statistical significance was evaluated using two tailed unpaired two sample equal variance Student’s t-test, with 0.05 threshold chosen for statistical significance; p-values indicated by #<0.1, * < 0.05, ** < 0.01, *** < 0.001.

## Data availability

All data are contained within the article. All raw data pertaining to the manuscript can be shared upon request to the corresponding authors.

## Acknowledgments

We thank Ana Farinho (Histology Facility, CEDOC, NMS-UNL) for Oil Red O staining and preparation of slides for microscopy. This research was funded to M. Paula Macedo by “Fundação para a Ciência e a Tecnologia” FCT (PTDC/BIM-MET/2115/2014); iNOVA4Health (UIDB/Multi/04462/2020); and by the Portuguese Diabetology Society.

## Author contributions

I.G.M. and M.P.M designed research; I.G.M. performed research; I.G.M. analyzed data; and I.G.M. and M.P.M wrote the paper.

